# tRNA-derived fragments restrict KSHV lytic replication through miRNA-like repression of host pro-viral signalling networks

**DOI:** 10.64898/2026.10.26.761788

**Authors:** Lorraine T. Matandirotya, James C. Murphy, Jessica M. Tucker, Adrian Whitehouse

**Affiliations:** School of Molecular and Cellular Biology, Faculty of Health and Life Sciences, University of Leeds, Leeds, LS2 9JT, UK; Astbury Centre for Structural Molecular Biology, University of Leeds, Leeds, LS2 9JT, UK; Department of Microbiology and Immunology, Carver College of Medicine, University of Iowa, USA

**Keywords:** KSHV, tRFs, RNA silencing, SHF

## Abstract

Transfer RNA-derived fragments (tRFs) are emerging regulators of gene expression. Despite mounting evidence of their functional relevance in cellular and pathological processes, their roles in virus infection remain poorly understood. Here, we show that host tRFs are dynamically reprogrammed during Kaposi’s sarcoma-associated herpesvirus (KSHV) lytic replication and function as potent antiviral effectors. Transcriptome-wide profiling revealed extensive remodelling of the tRF landscape following viral reactivation, identifying a subset of 5ʹ tRFs (tRF-His-GTG, tRF-Gly-CCC, and tRF-Lys-CTT) that are robustly induced independently of parental tRNA abundance. Functional analyses demonstrate that these tRFs suppress viral gene expression and infectious virion production. Mechanistically, we show that tRF-His-GTG and tRF-Lys-CTT engage canonical RNA silencing pathways by associating with AGO2 complexes and mediating sequence-specific repression of host target genes. Among these targets, we identify SHF as a critical pro-viral factor that promotes PI3K/Akt signalling, cellular survival and efficient viral replication. Collectively, our findings define a previously uncharacterised host defence pathway in which tRFs restrict KSHV replication by targeting host factors required for efficient viral propagation, thus expanding the functional repertoire of non-coding RNAs in antiviral immunity.

## Introduction

Viruses have evolved sophisticated strategies to evade host immune defences and enhance their replication. They encode their own repertoire of non-coding RNAs (ncRNAs) and also exploit host ncRNA networks to remodel the intracellular environment in favour of viral propagation^1–3^. Concurrently, several ncRNA species act as key regulators of immune responses, making them indispensable in cellular defence mechanisms. Consequently, ncRNAs represent a central interface in host-virus interactions, with both host and virus leveraging these molecules to shape infection outcomes^2^.

Kaposi’s sarcoma-associated herpesvirus (KSHV) is the causative agent of Kaposi’s sarcoma, primary effusion lymphoma, and multicentric Castleman disease^4–7^. A defining feature of KSHV biology is its biphasic lifecycle, consisting of latent persistence and lytic replication. Latency enables long-term maintenance of the viral genome, whereas lytic replication is essential for viral dissemination and pathogenesis^8,9^. KSHV is particularly adept at manipulating ncRNA-mediated regulatory pathways to influence host cellular processes during both phases of its life cycle. Specifically, virally-encoded miRNAs, circRNAs, and lncRNAs, together with virally-dysregulated host cell ncRNAs, have been shown to contribute to the establishment of latency, immune evasion, and control of the latent-to-lytic switch^10–15^.

Despite emerging evidence of their functional relevance, transfer RNA-derived fragments (tRFs) remain comparatively understudied relative to other ncRNA species. Once considered as generic by-products of tRNA degradation, it is now thought that tRFs are generated through the precise cleavage of precursor or mature tRNAs, and are classified according to their origin within the parental strands^16,17^. tRFs are gaining recognition as active regulators of gene expression, capable of mediating Argonaute (AGO)-dependent silencing, modulating RNA stability, inhibiting translation, and interacting with RNA-binding proteins^16,18–20^. These key features establish tRFs as potential modulators of both host and viral gene expression. Not surprisingly, tRFs are now being implicated in viral pathogenesis; either promoting or restricting virus replication in a context-specific manner. For instance, a proline-derived tRF-3 facilitates human T cell leukaemia virus type 1 replication by priming viral genomic synthesis, whilst respiratory syncytial virus utilises tRNA-GluCTC-derived tRF-5 molecules with miRNA-like activity to repress host targets and enhance viral replication^21,22^. Nevertheless, research into the roles of tRFs during viral infection remains in its infancy, with their potential roles in DNA virus infection largely unexplored. Interestingly, studies in murine gammaherpesvirus 68 (MHV68) have identified viral-derived tRF-like fragments, although their functional relevance is not yet understood^23^.

Here, we systematically investigate the role of cellular tRFs during KSHV lytic replication. We show that viral reactivation induces widespread remodelling of the tRF landscape and identify a subset of 5ʹ tRFs that are upregulated independently of their parental tRNA abundance. Functional analyses demonstrate that these tRFs act as potent inhibitors of viral replication, suppressing viral gene expression and the generation of infectious virions. Mechanistically, selected tRFs associate with AGO2 complexes and mediate sequence-specific repression of host target genes in a miRNA-like manner. Among these targets, we identify Src homology 2 domain-containing F (SHF) as a pro-viral factor that supports pathways linked to cell survival and viral fitness. Collectively, these findings not only establish tRFs as key components of the host antiviral response but also uncover a previously unrecognised layer of ncRNA-mediated regulation in KSHV infection.

## Results

### KSHV lytic reactivation induces widespread host tRF dysregulation

To determine whether cellular tRFs are dysregulated during KSHV infection, we analysed global tRF expression profiles by tRF-seq following lytic induction in TREx-BCBL1-RTA cells. Comparative analysis between latent and lytic conditions revealed extensive remodelling of the tRF landscape. Of the 574 tRFs identified, more than 300 tRF species showed significant changes in abundance, with 157 upregulated and 152 downregulated, respectively (**Figure 1A and B**). These findings demonstrate that KSHV lytic replication is associated with broad reprogramming of cellular tRF levels. Notably, a subset of tRFs exhibited particularly robust and consistent upregulation. From these, three 5ʹ tRFs derived from tRNA-His-GTG (tDR-1:32-His-GTG-1), tRNA-Gly-CCC (tDR-1:33-Gly-CCC-2), and tRNA-Lys-CTT (tDR-1:34-Lys-CTTCTT-1-M2) were selected for further investigation. These fragments, hereafter referred to as tRF-His-GTG, tRF-Gly-CCC, and tRF-Lys-CTT, represent discrete tRF isoforms generated from the 5’ of their parent tRNAs (Figure S1).

**Figure 1:**
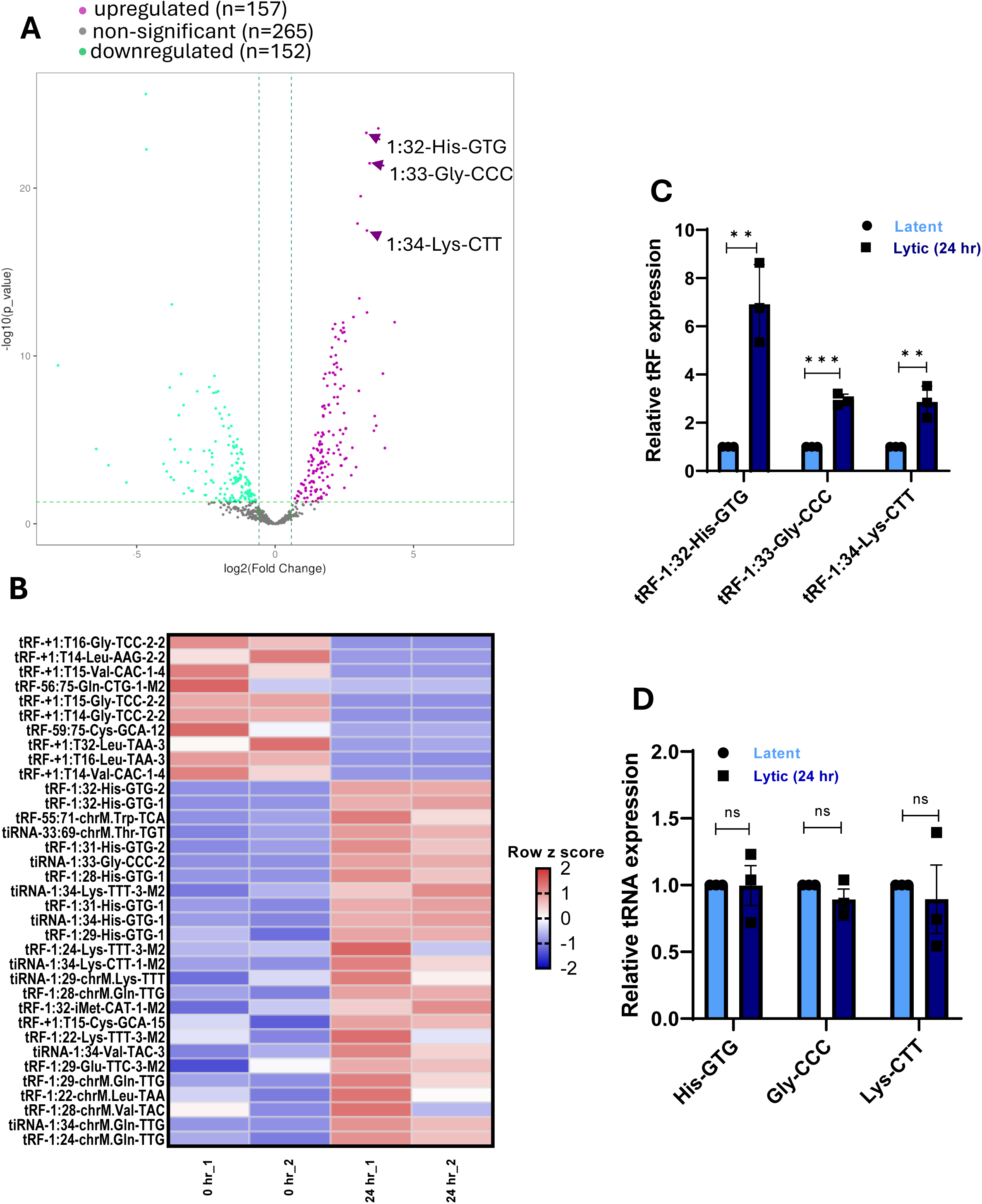
Cellular tRFs are dysregulated during KSHV lytic replication. (A and B) tRF-seq analysis of TREx BCBL1-RTA cells under latent conditions (0 h) and 24 h after KSHV lytic induction (n = 2). (A) Volcano plot of differentially expressed tRFs. (B) Heat map of upregulated and downregulated tRFs. (C) Expression of tRF-His-GTG, tRF-Gly-CCC and tRF-Lys-CTT latent and lytic (24 h) TREx-BCBL1-RTA cells, determined by RT-qPCR and normalised to *SNORD44*. (D) RT-qPCR analysis of the corresponding mature tRNAs shown in (C). Data in (C) and (D) are mean ± SD (n = 3). Statistical significance determined by unpaired two-tailed Student’s t-test (*p < 0.05, **p < 0.01, ***p < 0.001, and ****p < 0.0001).

To validate these findings, we employed a modified LNA-based RT-qPCR approach specifically optimised for tRF detection. This enabled selective amplification of short RNA fragments while discriminating against highly abundant parental tRNAs. We confirmed tRF-His-GTG, tRF-Gly-CCC, and tRF-Lys-CTT are significantly upregulated following the induction of KSHV lytic replication (**Figure 1C**). To determine whether this increase reflects changes in precursor availability, we examined the abundance of the corresponding mature tRNAs under the same conditions. Notably, no significant changes in tRNA levels were observed (**Figure 1D**), indicating that the increase in tRF abundance is not simply a consequence of increased expression levels within parental tRNA populations, but instead reflects regulated cleavage or processing events on a minor subpopulation of tRNAs. Together, these findings suggest tRF biogenesis is actively modulated during KSHV infection.

### Upregulated tRFs suppress KSHV gene expression and viral replication

Having established that specific tRFs are induced during KSHV lytic replication, we next determined their functional relevance. Based on the propensity of KSHV to reprogramme cellular ncRNAs in favour of viral outcomes, we hypothesised that tRF upregulation would enhance KSHV lytic replication. To test this, synthetic mimics of tRF-His-GTG, tRF-Gly-CCC, and tRF-Lys-CTT were transfected in TREx-BCBL1-RTA cells prior to lytic induction, and viral transcript levels were assessed (**Figure 2**). Unexpectedly, overexpression of each tRF resulted in a consistent and significant reduction viral transcripts. Specifically, quantitative RT-PCR analysis revealed that transcripts corresponding to an early viral gene (*ORF57*) showed comparatively minor changes, whereas later transcript levels, such as *ORF46* and *ORF26*, were more sensitive to tRF overexpression, suggesting that tRF-mediated effects may preferentially impact later stages of the lytic cascade (**Figure 2A-C**). To assess whether these transcriptional changes were reflected at the protein level, we performed immunoblot analysis of representative early and late viral proteins ORF57 and ORF26, respectively. Strikingly, tRF overexpression resulted in a pronounced reduction in protein abundance, exceeding the magnitude of change observed at the transcript level (**Figure 2D-F**). This suggests that tRFs exert their inhibitory effects on viral replication primarily at the level of protein production, and that this ultimately blocks the completion of the viral replication cycle. To confirm this, we assessed whether the effect of tRFs also impacted infectious virus production. TREx-BCBL1-RTA cells were transfected with tRF mimics or a negative control (NC) mimic and reactivated for 72 h with a high concentration of doxycycline (3 µg/μl), with latent cells included as an additional control. Cell-free supernatants were harvested and used to infect naïve HEK293T cells for 24 h. Viral genome load was then assessed in recipient HEK293T cells by qPCR of the ORF57 gene. Importantly, a negligible viral load was detected in latent controls, indicating that the transient transfection of synthetic mimics did not induce spontaneous lytic reactivation. Conversely, transfection of tRF-His-GTG, tRF-Gly-CCC, or tRF-Lys-CTT significantly reduced the production of infectious viral progeny (**Figure 2G**).

**Figure 2:**
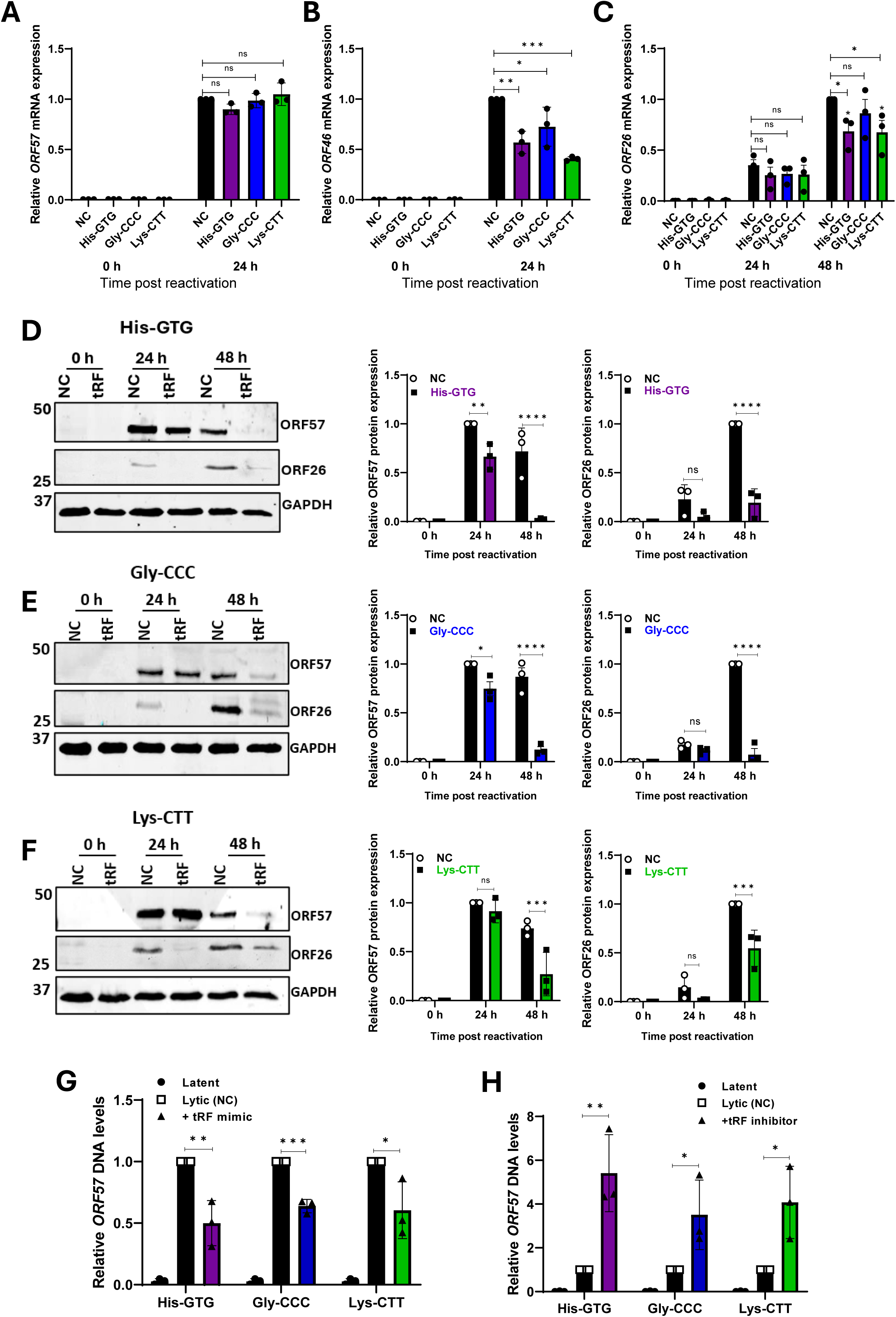
tRFs inhibit KSHV lytic replication. (A-F) TREx-BCBL1-RTA cells were transfected with control or tRF mimics before lytic induction. Samples were harvested at 0, 24 and 48 h post-induction. RT-qPCR analysis of (A) *ORF57*, (B) *ORF4C*, and (C) *ORF2C* transcripts normalised to *GAPDH*. (D-F) Immunoblot analysis of ORF57 and ORF26 protein levels following overexpression of tRF-His-GTG, tRF-Gly-CCC and tRF-Lys-CTT, with densitometric quantification normalised to GAPDH. (G-H) Viral re-infection assay. Following transfection of TREx-BCBL1-RTA cells with tRF mimics, antisense inhibitors, or respective controls prior to reactivation. Viral genome load was quantified by qPCR analysis of *ORF57* DNA normalised to *GAPDH*. (G) tRF mimic overexpression. (H) tRF inhibitor treatment. Data are mean ± SD (n = 3). Statistical significance determined by one-way ANOVA (A-C) or unpaired, two-tailed Student’s t test (D-H), (*p < 0.05, **p < 0.01, ***p < 0.001, and ****p < 0.0001).

To confirm tRF inhibition had an opposing effect on viral replication, LNA-enhanced antisense oligonucleotides (ASOs) were used to selectively perturb tRF function. In contrast to tRF overexpression, ASO-mediated inhibition significantly increased viral gene expression and enhanced virion output (**Figures 2H and S2**), although protein levels remained comparatively unchanged (**Figure S2**). Together, these findings establish tRFs as negative regulators of KSHV lytic replication, suppressing multiple stages of the viral life cycle.

### tRF-His-GTG and tRF-Lys-CTT function through miRNA-like mechanisms

To define the molecular mechanisms underlying tRF-mediated antiviral activity, we next examined the subcellular localisation and protein associations of tRF-His-GTG, tRF-Gly-CCC, and tRF-Lys-CTT. As RNA localisation often provides insight into regulatory function, we first assessed the intracellular distribution of these tRFs in TREx-BCBL1-RTA cells using biotin-labelled synthetic mimics. Under both latent and lytic conditions, the biotinylated-control resulted in a diffuse distribution throughout cytoplasmic and nuclear compartments. Conversely, all three tRFs primarily exhibited a cytoplasmic localisation (**Figure 3A**), suggesting potential roles in post-transcriptional regulation rather than nuclear processes such as transcriptional control or chromatin modification. Notably, each tRF led to the formation of discrete puncta within the cytoplasm, consistent with the potential association with ribonucleoprotein complexes. Nevertheless, it was important to rule out the possibility that the observed cytoplasmic tRF localisation was an artefact from the transient transfection of exogenous mimics. Therefore, the cytoplasmic localisation of native tRFs was confirmed by subcellular fractionation and subsequent RT-qPCR analysis (**Figure 3B**).

**Figure 3:**
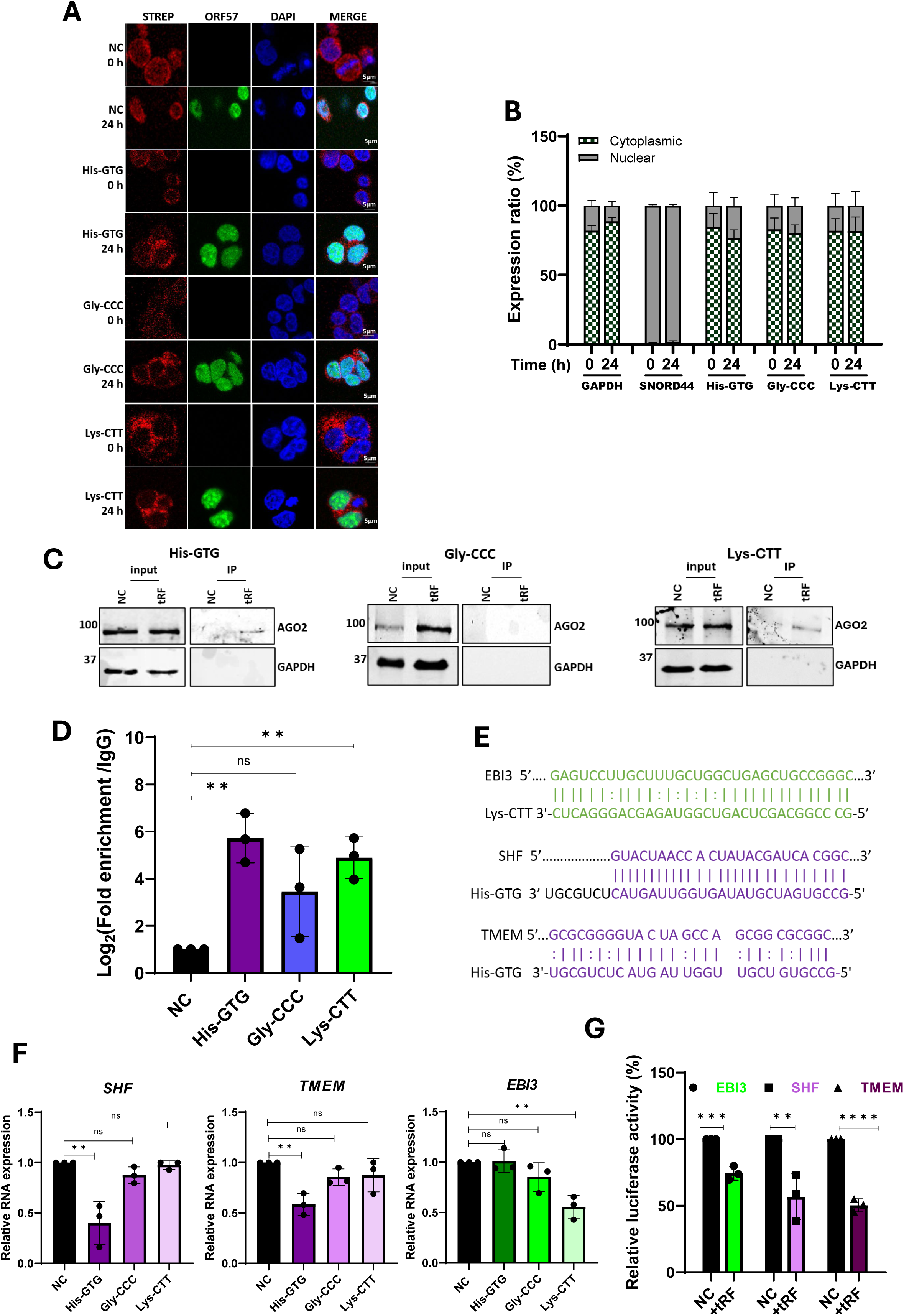
tRFs His-GTG and Lys-CTT function through miRNA-like mechanisms. (A) Subcellular distribution of biotinylated-tRF mimics or controls in TREx-BCBL1-RTA cells during latency and 24 h post reactivation. Cells were stained for streptavidin, ORF57 and DAPI. Scale bars represent 5 µm. (B) Cytoplasmic and nuclear distribution of endogenous of tRF-His GTG, tRF-Gly CCC and tRF-Lys-CTT, as assessed by RT-qPCR. *GAPDH* and *SNORD44* served as fractionation controls. Data presented as percentage of total cellular tRF abundance. (C) Streptavidin pull-down assay in TREx-BCBL1-RTA cells lysates expressing biotinylated-control or tRF mimics, followed by immunoblotting for AGO2 and GAPDH. (D) AGO2 RNA immunoprecipitation followed by RT-qPCR analysis of associated tRFs. IgG served as a negative control, input RNA was normalised to *SNORD44*. (E) Predicted tRF-traget interactions, using *tRFtarget*. (F) RT-qPCR analysis of predicted targets (*SHF*, *TMEM*, and *EBI3*) following tRF overexpression in TREx-BCBL1-RTA cells, normalised to *GAPDH*. (G) Dual-luciferase reporter assays in HEK293T cells co-transfected with SHF or TMEM 5′UTR, or EBI3 3′UTR reporter constructs and indicated tRF mimics. Luciferase activity was normalised to the negative control. Data are mean ± SD (n = 3). Statistical significance determined by one-way ANOVA (D and F) or unpaired, two-tailed Student’s t test (G), (*p < 0.05, **p < 0.01, ***p < 0.001, and ****p < 0.0001).

To investigate the molecular mechanisms governing tRF functionality, we next assessed whether tRF-His-GTG, tRF-Gly-CCC and tRF-Lys-CTT associate with Argonaute proteins and exhibit miRNA-like activity, previously reported for some tRFs^18,24–26^. Streptavidin pull-down assays using biotinylated-tRF mimics demonstrated that tRF-His-GTG and tRF-Lys-CTT interact with AGO2, whereas tRF-Gly-CCC failed to show any detectable enrichment (**Figure 3C**). These findings were further validated using reciprocal RNA immunoprecipitation assays, confirming selective incorporation of tRF-His-GTG and tRF-Lys-CTT into AGO2-containing complexes (**Figure 3D**).

We next assessed whether these interactions confer sequence-specific gene silencing capability. Computational target prediction, using the tRFtarget prediction platform^27^, identified candidate mRNAs with potential binding sites within their untranslated regions. Although tRF-Gly-CCC was not enriched in AGO2 complexes, it was included in the target screen to account for potential AGO2-independent gene regulatory mechanisms. The putative targets with highest confidence, based on free energy and complementary sequence length, are summarised in **Table S1**. Among these, Src homology 2 domain-containing F (SHF) and transmembrane protein 237 (TMEM237, hereafter referred to as TMEM) were prioritised as candidate targets of tRF-His-GTG, while Epstein-Barr virus-induced gene 3 (EBI3) was identified as a candidate target of tRF-Lys-CTT (**Figure 3E**). Interestingly, all three participate in signalling networks that are dysregulated in diseased states^28–31^.

The putative targets were subsequently validated by assessing their expression in TREx-BCBL1-RTA cells in the presence of tRF mimics (**Figure S3**). Overexpressing tRF-His-GTG significantly reduced *SHF* and *TMEM* transcript levels, whereas overexpressing tRF-Lys-CTT suppressed *EBI3* (**Figure 3F**). Importantly, only cognate tRF overexpression depleted transcript levels, whereas transfection of non-targeting mimics produced comparable expression profiles with the negative control, suggesting functional mechanisms involving targeted regulatory interactions rather than global transcriptional repression. In contrast, none of the predicted tRF-Gly-CCC targets showed altered expression upon tRF-Gly-CCC overexpression (**Figure S3**), consistent with its lack of AGO2 interaction.

Next, direct sequence-mediated repression by tRF-His-GTG and tRF-Lys-CTT was confirmed using dual-luciferase reporter assays. HEK293T cells were co-transfected with reporter constructs containing 5’ UTRs of SHF or TMEM; or the 3’UTR of EBI3, along with the corresponding tRF mimic or NC mimic. In each instance, insertion of predicted binding sites into reporter constructs conferred sensitivity to tRF-mediated repression (**Figure 3G**). Collectively, these findings demonstrate that tRF-His-GTG and tRF-Lys-CTT engage canonical RNA silencing machinery to mediate sequence-specific repression of host transcripts and potentially link tRF function directly to antiviral roles.

### tRF-mediated repression targets pro-viral host factors during KSHV infection

Considering the inhibitory effects of tRF-His-GTG and tRF-Lys-CTT on KSHV replication, we next explored the possibility that their target genes function as pro-viral host factors. Analysing the expression dynamics of SHF, TMEM and EBI3 over a 48 h period following reactivation, showed all three targets exhibited a discordant pattern between RNA and protein levels. Transcript levels of *SHF*, *TMEM*, and *EBI3* increased transiently during early stages of lytic reactivation, whereas protein levels declined progressively (**Figure S4**). This observation suggests that post-transcriptional regulatory mechanisms override the transcriptional induction visible at 24 h post-reactivation (**Fig S4A**), consistent with AGO2-mediated repression. To disentangle the opposing effects of viral lytic induction from tRF activity, we modulated tRF levels under latent and lytic conditions and monitored target gene expression. Overexpression of tRF-His-GTG or tRF-Lys-CTT reduced target gene expression (**Figure 4A and B**), whereas tRF inhibition restored or enhanced target expression (**Figure 4C and D**). Notably, effects were most pronounced under lytic conditions, where target protein expression was almost completely abrogated in the presence of tRF mimics, compared to control samples. This amplified antiviral phenotype reflects the potential outcome of coupling intrinsic, upregulated tRF activity during lytic reactivation, with the repression imposed by overexpressing synthetic mimics. Overall, these findings indicate that tRF-mediated repression represents a regulatory mechanism controlling the expression of specific host factors during infection.

**Figure 4:**
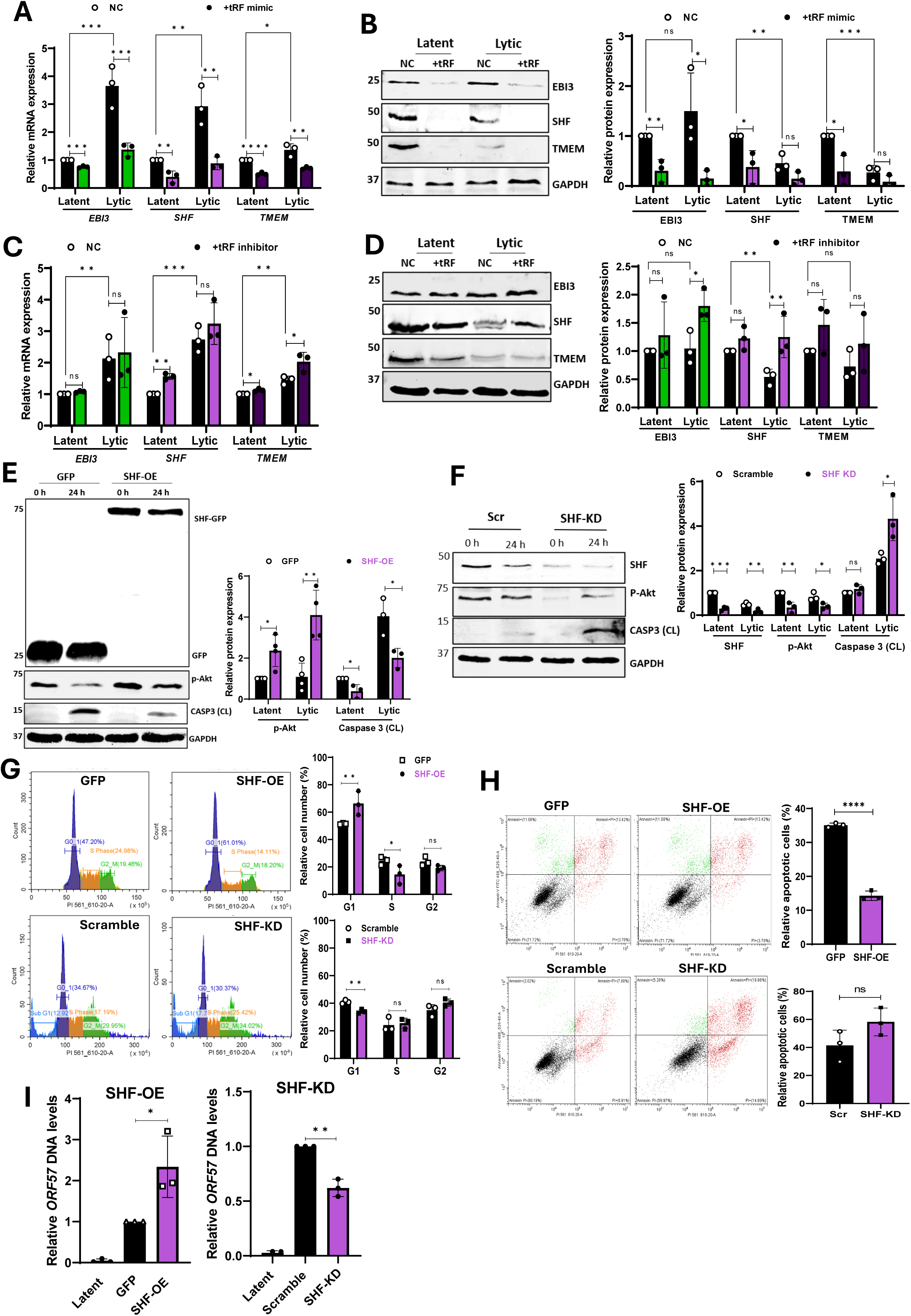
tRF-mediated repression targets pro-viral host factors during KSHV infection. (A-D) TREx-BCBL1-RTA cells transfected with tRF mimics, antisense inhibitors, or respective controls were analysed during latency and 24 h post lytic-induction. (A) RT-qPCR analysis of *EBI3*, *SHF*, and *TMEM* expression following tRF overexpression, normalised to *GAPDH*. (B) Corresponding immunoblot analysis from (A), with densitometric quantification normalised to GAPDH. (C) RT-qPCR analysis of *EBI3*, *SHF*, and *TMEM* expression following tRF inhibition, normalised to *GAPDH*. (D) Corresponding immunoblot analysis from (C), with densitometric quantification normalised to GAPDH. (E) Immunoblot analysis of TREx-BCBL1-RTA cell lysates stably expressing GFP-SHF (SHF-OE) or GFP-control at 0 h and 24 h post reactivation, probed for GFP, p-Akt, cleaved Caspase 3 and GAPDH. Densitometric analysis normalised to GAPDH. (F) Immunoblot analysis TREx-BCBL1-RTA cell lysates stably expressing shRNAs targeting SHF (SHF-KD) and scramble controls at 0 h and 24 h post reactivation, probed for SHF, p-Akt, Caspase 3 and GAPDH. Densitometric analysis normalised to GAPDH. (G) Cell cycle analysis of SHF-OE, SHF-KD, and control cells 24 h post reactivation using propidium iodide staining, with quantification of G1, S and G2 populations. (H) Annexin V-FITC/PI apoptosis analysis of SHF-OE, SHF-KD, and control cells 24 h post-induction, with quantification of apoptotic cells. (I) Viral re-infection assays using SHF-OE and SHF-KD cells. Infectious virion production was quantified by qPCR analysis of *ORF57* DNA normalised to *GAPDH*. Data are mean ± SD (n = 3). Statistical significance determined by one-way ANOVA (A, B and G) or unpaired, two-tailed Student’s t test (C-F and H-L), (*p < 0.05, **p < 0.01, ***p < 0.001, and ****p < 0.0001).

### SHF functions as a pro-viral host factor that promotes KSHV replication

Next, we investigated the pro-viral activity of tRF target genes by assessing their ability to enhance KSHV replication. TREx-BCBL1-RTA were stably transduced with lentivirus-based gene-targeting shRNAs or scramble controls, and loss-of-function studies performed. Depletion of either EBI3, SHF or TMEM all resulted in reduced expression of key viral genes, particularly those associated with later stages of lytic replication; as previously observed with tRF mimic overexpression studies, indicating that these host factors support efficient viral replication (**Figure S4**).

SHF consistently displayed the most pronounced phenotype and was therefore selected for further characterisation. SHF is an adaptor protein implicated in multiple signalling pathways, including PI3K/Akt and JAK/STAT signalling, known to be exploited during KSHV infection^31–33^. Therefore, stable TREx-BCBL1-RTA cell lines overexpressing GFP-SHF (SHF-OE) or GFP alone were generated and the impact of SHF overexpression on the levels of downstream cytokines was assessed under both latent and lytic conditions. Overexpressing SHF led to enhanced levels of phosphorylated Akt (p-Akt), with a corresponding reduction in cleaved caspase 3 levels, indicating suppression of apoptosis-related cellular responses (**Figure 4E**). Conversely, SHF depletion resulted in reduced p-Akt levels and increased cleaved caspase 3 during lytic replication, indicating reciprocal effects on these signalling pathways (**Figure 4F**).

The manipulation of host cell cycle progression is among many exploitative strategies employed by KSHV. By halting cells in G1 phase, KSHV promotes a stable environment for viral DNA synthesis whilst simultaneously avoiding competition for resources with host genome replication in S phase^1^. Moreover, delaying cell cycle progression effectively stalls cell cycle checkpoints that would otherwise trigger premature apoptosis prior to the successful packaging and release of infectious progeny^32,34,35^. Following the effects of SHF dysregulation on Akt signalling, we investigated whether SHF influenced cell cycle progression (**Figure 4G**). Overexpression of SHF increased the proportion of cells in G1 phase, while reducing S phase progression. Conversely, SHF depletion disrupted this balance and increased the proportion of apoptotic cells. Consistent with the observed effects on cell cycle distribution, Annexin V-FITC staining revealed a significant reduction in apoptosis with SHF overexpression, whereas knockdown produced modest increases in cellular apoptosis, further highlighting a role of SHF in pro-survival pathways during lytic replication (**Figure 4H**).

These data suggest that SHF contributes to the establishment of a cellular environment supporting efficient viral replication by promoting survival signalling, limiting apoptosis, and modulating cell cycle progression. Concordant with the proposed pro-viral role of SHF, overexpression was associated with a significant increase in infectious progeny, whereas SHF knockdown effectively decreased infectious virion levels (**Figure 4I**). Notably, these findings closely recapitulate the effects of tRF-His-GTG dysregulation on viral fitness, validating a functional relationship between tRF-mediated SHF repression and the regulation of KSHV lytic replication.

## Discussion

This study identifies transfer RNA-derived fragments (tRFs) as regulators of KSHV lytic replication and demonstrates that they function as antiviral effectors through the targeted suppression of host factors. These findings expand our understanding of non-coding RNA-mediated host-virus interactions and highlight tRFs as previously underappreciated components of antiviral defence. We show that tRF-His-GTG, tRF-Gly-CCC, and tRF-Lys-CTT are selectively upregulated during KSHV lytic replication and suppress viral gene expression, protein production, and infectious virion output. Importantly, these effects occur independently of parental tRNA abundance, suggesting that KSHV infection alters tRNA processing, consistent with observations in other gammaherpesviruses^36,37^.

Mechanistically, our data demonstrate that tRF-His-GTG and tRF-Lys-CTT tRFs exhibit miRNA-like activity. Both localise to the cytoplasm, associate with AGO2, and mediate sequence-specific repression of target mRNAs, supporting growing evidence that subsets of tRFs utilise canonical RNA silencing pathways^26,38,39^. In contrast, tRF-Gly-CCC neither interacted with AGO2 nor suppressed candidate targets, highlighting functional and mechanistic diversity among tRF species.

A major finding of this study is the identification of SHF as a novel pro-viral host factor targeted by tRF-His-GTG. SHF plays a central role in regulating signalling pathways that are critical for cellular survival, proliferation, and metabolism^31,33^, pathways frequently co-opted by KSHV to promote efficient replication and evade host defences^32^. SHF enhances Akt signalling, reduces apoptosis, and promotes G1 phase cell cycle arrest, thereby creating a cellular environment favourable for viral replication. Repression of SHF by tRF-His-GTG therefore represents a targeted antiviral strategy whereby host cells limit viral propagation by disrupting signalling pathways exploited by the virus. This aligns with a broader model of host-virus conflict, in which viruses induce the expression of host factors that facilitate replication while the host simultaneously activates mechanisms to counteract these effects. he observed increase in SHF transcript abundance despite reduced protein levels further suggest competing regulatory pressures between host antiviral responses and viral replication programmes.

Beyond SHF, we also identify TMEM and EBI3 as cellular targets of tRF-mediated repression. Although their roles in KSHV replication were not fully characterised, both are linked to pathways involved in oncogenesis and immune regulation^28,40,41^. Future studies will be required to determine whether these cellular targets play complementary roles in supporting viral replication and whether their repression contributes to antiviral outcomes. While our study focused primarily on host targets, the ability of tRFs to engage AGO2 and mediate sequence-specific repression raises the possibility that tRFs may also regulate viral transcripts directly. Such interactions could provide an additional layer of antiviral regulation, however, an alternative model is that they primarily target host dependency factors, providing a broader antiviral response that is less vulnerable to viral sequence variation. However, the extensive tRF dysregulation observed during infection further suggests that distinct tRFs may exert both antiviral and pro-viral functions.

More broadly, our findings reinforce the emerging view of tRFs as versatile regulators of gene expression with context-dependent functions. The ability to engage both canonical silencing pathways and potentially alternative mechanisms underscores their adaptability to respond dynamically to changing cellular conditions and exert fine-tuned control over both host and viral processes. Notably, because herpesviruses often suppress RNA polymerase II-dependent transcription while maintaining or enhancing RNA polymerase III-driven tRNA transcription, tRF-mediated may provide a robust antiviral mechanism when host miRNA biogenesis is impaired^42,43^.

In conclusion, we establish tRFs as previously unrecognised components of the host antiviral response to KSHV infection. By repressing host factors required for viral replication, tRFs restrict lytic replication and viral propagation. These findings broaden our understanding of ncRNA biology and highlight tRFs as potential therapeutic targets in KSHV-associated diseases.

### Limitations of the study

Although this study clearly establishes the induction of functional tRFs during KSHV lytic replication, the mechanisms governing their biogenesis remain unclear. Endonucleases, such as Angiogenin and Dicer, have been implicated in tRF biogenesis, however their relative contributions remain unclear^44,45^. Defining how tRFs accumulate during lytic infection therefore represents an important avenue for future research. Additionally, given that synthetic tRF mimics lack endogenous RNA modifications, the influence of tRNA modifications on tRF biogenesis and activity remains unknown.

Mechanistic studies focused primarily on SHF, while the contributions of EBI3 and TMEM to KSHV lytic replication were only partially explored. Given the roles of EBI3 in herpesvirus-associated immune regulation and TMEM in WNT/ β-catenin signalling^40,46^, future work should determine whether these host factors similarly facilitate KSHV replication.

Finally, the possibility that KSHV generates specialised tRF-like molecules also warrants further investigation. KSHV extensively remodels the host epitranscriptome^47–50^, and RNA modifications are known to impact both tRF biogenesis and function^51–53^. Viral tRFs (virtRFs) have recently been identified in MHV68^36^, raising the possibility that KSHV may similarly generate distinct tRF species. Although we found no evidence for such molecules, the extensive viral reprogramming of cellular RNA-processing pathways, including specialised ribosomes and paraspeckles during lytic replication^49,54,55^, suggests that virus-derived or virus-modified tRFs could contribute to efficient KSHV replication.

## Acknowledgements

This work was supported in parts by the Medical Research Council to AW (MR/V009478/1 and MR/X000060/1), Wellcome Trust to AW (318573/Z/24/Z), BBSRC to LM and AW (BB/T007222/1).

## Author Contributions

Conceptualization (LM, AW); Data curation (LM); Formal Analysis (LM, JT, AW); Funding acquisition (AW); Investigation (LM, JM); Writing – original draft (LM, AW); Writing – review & editing (All authors).

## Material & Methods

## Key Resources Table

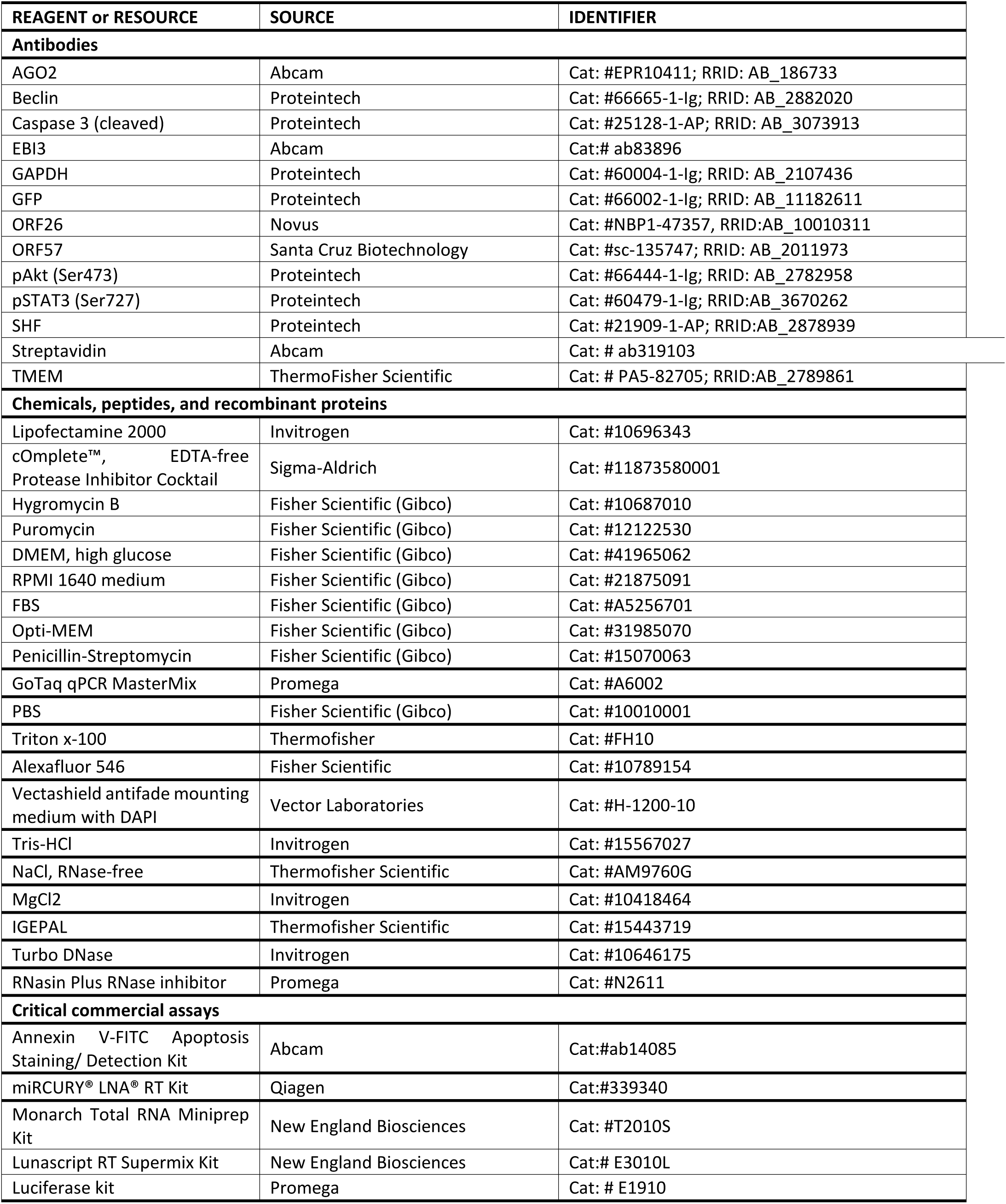
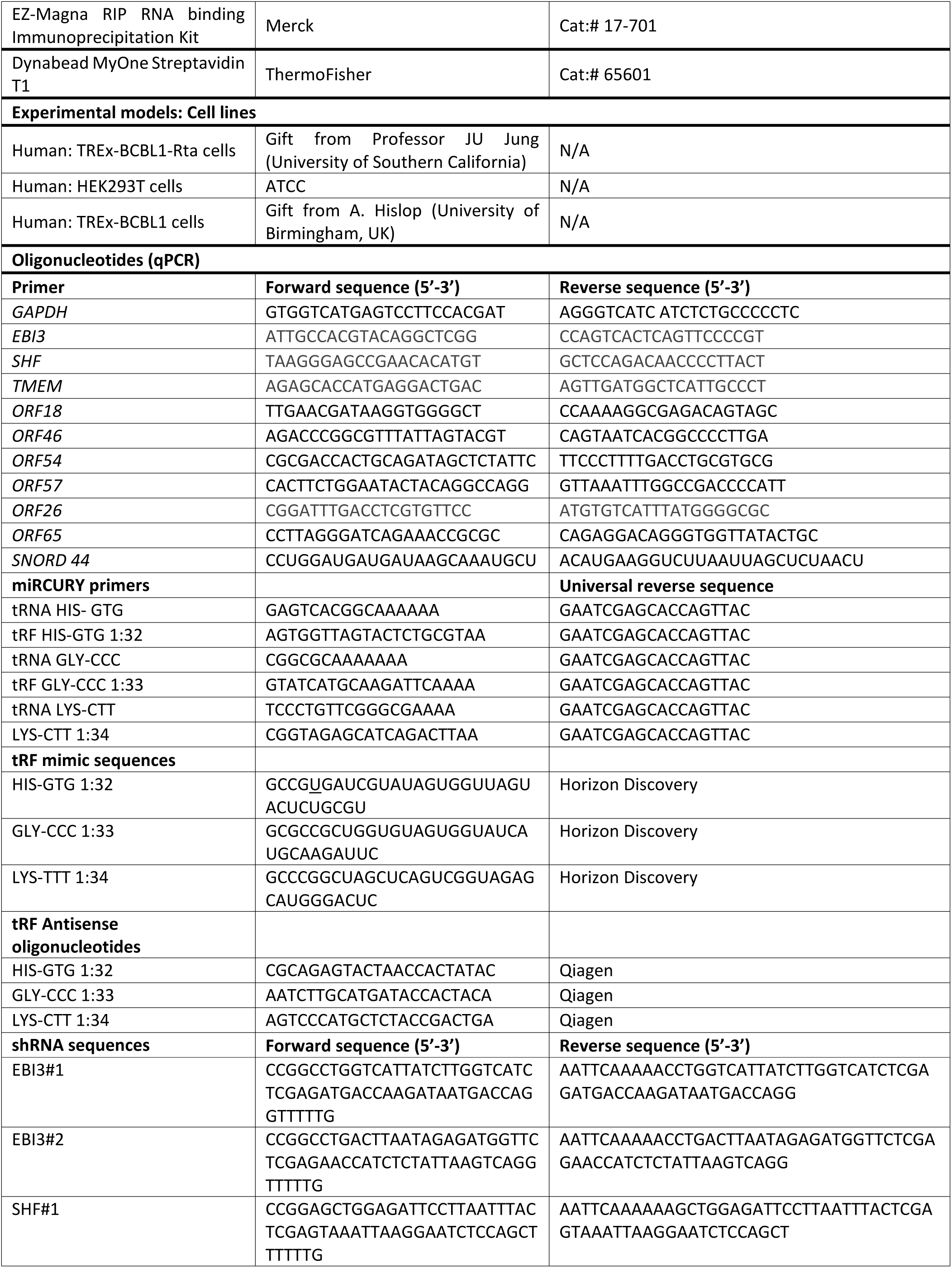
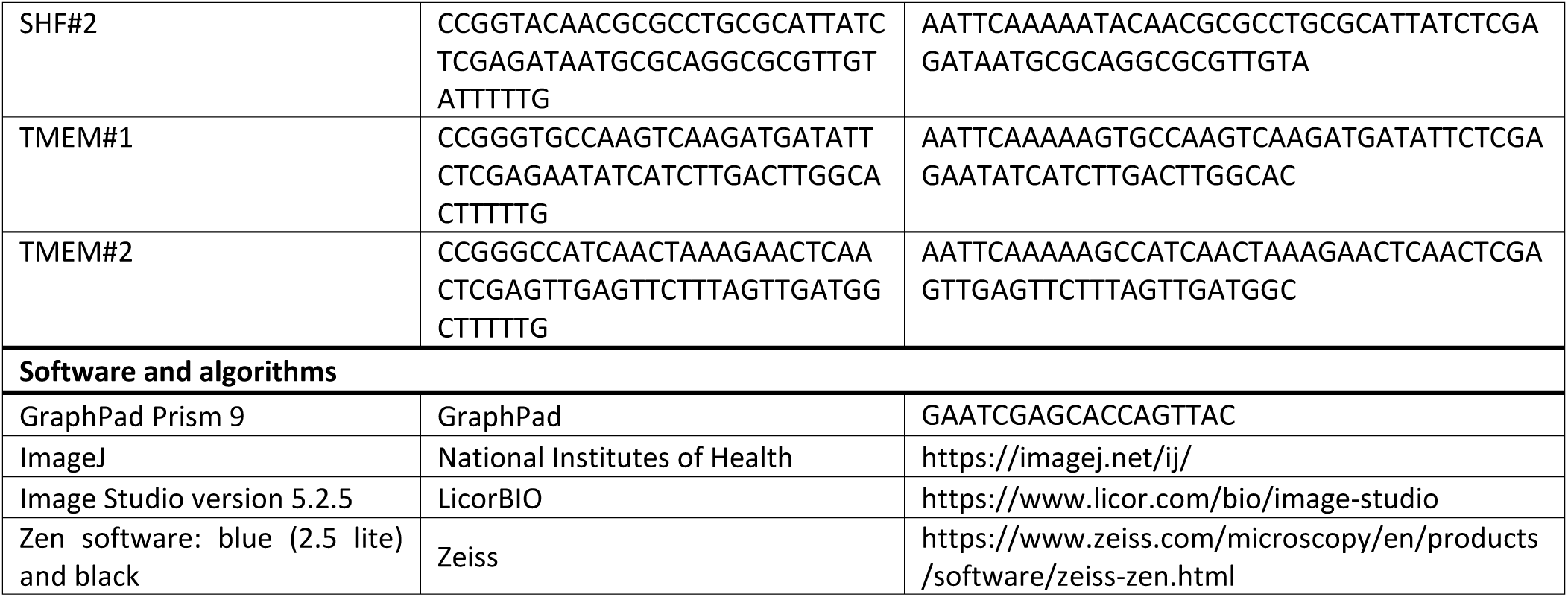

### Methods

#### Cell culture

TREx-BCBL1-RTA cells, a B cell lymphoma cell line latently infected with KSHV engineered to contain a doxycycline inducible myc-RTA were a gift from Professor JU Jung (University of Southern California). TREx-BCBL1-RTA cells were cultured in RPMI1640 with glutamine (Gibco), supplemented with 10% Foetal Bovine Serum (FBS) (Gibco), 1% P/S (Gibco) and 100 µg/ml hygromycin B (ThermoFisher). Unless stated otherwise, virus lytic replication in TREx-BCBL1-RTA cells was induced via addition of 2 µg/ml doxycycline hyclate (Sigma-Aldrich). HEK293T cells were purchased from the ATCC and cultured in DMEM (Lonza) and supplemented with 10% FBS and 1% P/S. All cell lines were tested negative for mycoplasma.

#### Plasmid constructs

The packaging plasmids pVSV.G and psPAX2 were a gift from Dr Edwin Chen (University of Westminster, London). psiCheck2 was a gift from Dr James Boyne (Leeds Beckett University). pLenti-CMV-GFP was a gift from Dr Ollie Manners (University of Leeds). pLKO.1 TRC cloning vector was purchased from Addgene (gift from David Root; Addgene plasmid #10878) with shRNAs with against EBI3, SHF and TMEM cloned into it. The scramble sequence is: CCGCAGGTATGCACGCGT.

### Method details

#### RNA extraction and qPCR

Total RNA was extracted using Monarch Total RNA Miniprep Kit (NEB) as per manufacturer’s protocol. 1 µg RNA was reverse transcribed using LunaScript RT SuperMix Kit (NEB). qPCR was performed using synthesised cDNA, GoTaq qPCR MasterMix (Promega) and the appropriate primer. qPCR was performed on Rotorgene Q and analysed by the ΔΔCT method against a housekeeping gene as previously described before plotting as fold change^56^. For quantification of tRFs, the miRCURY LNA miRNA RT kit and miRCURY LNA SYBR Green PCR Kits were used for reverse transcription and cDNA amplification, respectively, and instructions were followed as provided by the manufacturer.

#### tRF-Seq

tRF-Seq samples were processed by Arraystar Inc. Total RNA was extracted from TREx BCBL1-RTA cells at 0 and 24 h post lytic induction and pretreated as follows for efficient reverse transcription and small RNA-seq library construction: 3’-aminoacyl deacylation, 3’-cP removal, 5’-OH phosphorylation, and m1A and m3C demethylation. Samples were assessed using an Agilent High Sensitivity D1000 Screen Tape Station to ensure RNA integrity. Sequencing libraries were size-selected for the RNA biotypes using an automated gel cutter and an Agilent BioAnalyzer 2100. cDNA libraries were analysed by Illumina NextSeq 500 system as per manufacturer instructions. Bioinformatics analysis was performed by Arraystar Inc. Raw reads were processed for RNA expression using the Solexa pipeline v1.8 (Off-Line Base Caller software, v1.8). Quality filtered (FastQC) and trimmed reads (cutadapt) were aligned to mature tRNA sequences using Bowtie2 (v 2.4.2.), allowing for one mismatch only to precursor tRNAs. Expression levels were normalised by “counts per million” (CPM). Differential expression analyses between two KSHV replication time points were performed using the R package edgeR.

#### Transient transfection

NC and tRF mimics were designed and synthesized by Dharmacon Inc. Antisense oligonucleotides (ASOs) and NC inhibitors were designed and synthesised by Qiagen. TREx-BCBL1-RTA cells were transfected using the Lipofectamine RNAiMAX Transfection Reagent (Thermofisher) as per the manufacturer’s instructions. Briefly, cells were seeded at a density of 7.5×10^5^ cells/ml and inoculated with a final concentration of 50 nM tRF mimics or 100 nM ASOs as reported previously,^26^ unless stated otherwise. Prior to transfection, mimics and ASOs were pre-incubated for 20 min at room temperature (RT) with 4 µl RNAiMAx diluted in 500 µl of Opti-MEM Reduced Serum Medium. Samples were reactivated 4 h post transfection and harvested at 24 h or 48 h post lytic induction, as specified.

#### Immunoblotting

Cell lysates were separated using 8-12% polyacrylamide gels and transferred to Amersham Nitrocellulose Membranes (GE healthcare) via Trans-blot Turbo Transfer system (Bio-Rad). Membranes were blocked in TBS + 0.1% tween with 5% w/v dried skimmed milk powder. Membranes were probed with appropriate primary antibodies and Dylight800 and 600 secondary antibodies at 1:10000 dilution (ThermoFisher). Proteins were detected using the LICOR.

#### Viral reinfection assays

TREx-BCBL1-RTA cells were seeded at a density of 7.5×10^5^ cells/ml and reactivated with a high concentration of doxycycline (3 µg/ml) for 72 h, with one uninduced sample included per test. The supernatant, containing any released virion particles, was filtered through a 0.45 µm filter (Merck Millipore) and added in a 10:1 ratio with HEK293T growth medium onto HEK293T cells at 70% confluency. The cells were incubated for 24 h, then DNA was isolated from the harvested cell pellets. Purified DNA samples were analysed by qPCR for the levels of *ORF57* to determine infectious virion production as previously described^48^.

#### Immunofluorescence

TREx-BCBL-1-RTA cells were transfected with biotinylated tRF mimics 24 h prior to seeding onto poly-L-lysine (Sigma-Aldrich) coated coverslips and reactivated 3 h later. The cells were fixed in PBS containing 4% paraformaldehyde (v/v) for 15 min, then permeabilised in PBS containing 1% Triton X-100 for 15 min. All the following steps were incubated for 1 h at 37°C. Coverslips were blocked in PBS containing 1% BSA, incubated in anti-streptavidin (1:100), then incubated in Alexa Fluor 546 (1:500) before being mounted onto slides using Vectashield hardset mounting medium containing DAPI. Slides were visualised on a Zeiss LSM880 laser scanning confocal microscope and processed using ZEN imaging software.

#### Subcellular fractionation

A total of 1 × 10^6^ TREx-BCBL1-RTA cells were harvested per condition and lysed in 600 μl PBS with 1% (V/V) triton-X 100 and 1 µl/ml Murine RNase inhibitor at 4 °C for 10 min. A 250 µl fraction of the extract was taken as a whole cell lysate, while the remaining 350 µl was centrifuged at 720 x g for 5 min at 4°C. The supernatant was collected as the cytoplasmic fraction while the pelleted nuclei were washed in 1 ml PBS and centrifuged once again at 720 x g for 5 min at 4 °C. The pelleted nuclei were resuspended in 100 µl PBS (1% triton-X-100, 1 µl/ml Murine RNase inhibitor). The fractions were then deposited in TRIzol LS reagent for assessment of RNA levels.

#### RNA immunoprecipitations (RIPs)

TREx-BCBL1-RTA cells were seeded at a final density of 5 × 10^6^ cells per condition and RNA-protein complexes were fixed by UV crosslinking (150 mJ/cm²) at the time of cell harvesting. AGO2 RIPs were conducted using the EZ-Magna RIP RNA-binding Immunoprecipitation kit (Merck millipore) as per manufacturer’s instructions. RNA was extracted and purified using TRIzol LS (Invitrogen) as per manufacturer’s instructions before analysis via RT-qPCR. Samples were analysed using fold enrichment over percentage inputs.

For reciprocal immunoprecipitations, TREx-BCBL1-RTA cells were seeded at a final density of 5 × 10^6^ cells per condition and transiently transfected with biotinylated tRF mimics for 24 h. RNA-protein complexes were fixed by UV crosslinking (150 mJ/cm²) at the time of sample harvesting, and streptavidin pull downs were performed using Dynabead MyOne Streptavidin T1 (Thermofisher) beads as previously described^57^. Co-precipitated proteins were eluted from the beads by resuspension in 30µl 2x SDS sample buffer^58^, and tRF association with AGO2 complexes was subsequently determined by immunoblot analysis.

#### Luciferase assays

Luciferase activity was detected using the Dual-Luciferase Reporter Assay System (Promega). The 5’ or 3’ UTR of EBI3 (NCBI Gene ID: 10148), SHF (NCBI Gene ID: 90525) and TMEM (NCBI Gene ID: 65062) were cloned upstream of the renilla reporter gene in the psiCHECK™-2 vector dual-promoter construct, with a control firefly reporter gene on a separate promoter. HEK293T cell lines were seeded in triplicate at a density of 3 × 10^5^ cells per well and grown for 24 h. Cells were transfected with 100 ng of respective plasmids (diluted in 100 µL Opti-mem) using 3 µL Lipofectamine® 2000 (InvitrogenTM; diluted in 100 µL Opti-mem), in the presence of a NC mimic or each tRF mimic and incubated for a further 24 h. Media was removed from the wells and cells washed gently with 100 μl PBS. Aliquots (100 μL) of 1x passive lysis buffer were added to the cell monolayers and rocked for 20 min at room temperature and then 10 μL of each lysate was transferred to tissue culture treated white microplates (Greiner Bio-One). Luciferase measurements were carried out using a FLUOstar Optima microplate reader (BMG Labtech Ltd), with injectors 1 and 2 being used to dispense 50 μL of Luciferase Assay Reagent II and Stop & Glo Reagent respectively. Renilla luciferase activity was normalized to Firefly luciferase activity.

#### Lentivirus transduction

Lentiviruses were generated by transfection of HEK293T cells seeded in 12-well plates using a three-plasmid system as described previously^59^. Per 12-well, 4 µl of lipofectamine 2000 (Thermo Scientific) were used together with 1.2 µg of the appropriate lentiviral vector, 0.65 µg of pVSV.G, and 0.65 µg psPAX2. Eight hours post-transfection, media was changed with 1.5 mL of DMEM supplemented with 10% (v/v) FBS. Two days post-transfection, viral supernatants were harvested, filtered through a 0.45 µm filter (Merck Millipore) and immediately used for transductions of TREx BCBL1-RTA cells. Cells (5×10^5^) in 12 well plates were infected by spin inoculation for 60 min at 800 x g at room temperature, in the presence of 8 μg/ml of polybrene (Merck Millipore). Puromycin (3 µg/ml) was added 48 h post-transduction before stable knockdown/ overexpression was analysed via qPCR and western blot if appropriate.

#### Cell cycle analysis

SHF-GFP overexpressing or SHF-depleted TREx-BCBL1-RTA cells (along with relevant controls) were seeded at a density of 5 ×10^5^ cells/ml and reactivated for 24 h. Cells were harvested and fixed O/N at -20 °C in 70% ethanol. The fixed cells were washed twice by resuspending them in PBS (0.5% BSA) and centrifugation (500 x g for 10 min at 4 °C). Cells were stained for 30 min at RT in PBS (0.5% BSA), 5 μg/ml RNase and 25 μg/ml propidium iodide (Sigma). Samples were processed on CytoFlex S Benchtop Flow Cytometer (Beckman Coulter).

#### Apoptotic assays

SHF-GFP overexpressing or SHF-depleted TREx-BCBL1-RTA cells (along with relevant controls) were seeded at a density of 5 ×10^5^ cells/ml and reactivated for 24 h. Apoptotic events were detected using the Annexin V-FITC Apoptosis Staining/Detection kit (Abcam) as outlined by the manufacturer. In brief, cells were harvested and washed twice with PBS. The cells were then resuspended in 500 μl of 1x binding buffer and stained with 5 μl each of Annexin V-FITC and PI for 30 min at RT in the dark. Samples were processed on CytoFlex S Benchtop Flow Cytometer (Beckman Coulter).

#### Quantification and statistical analysis

All quantitative data are presented as the mean ± standard deviation (SD) from at least three biologically independent experiments unless stated otherwise. Comparisons between two independent groups were analysed using an unpaired two-tailed Student’s t-test. Where experiments involved comparisons between three or more groups, one-way analysis of variance (ANOVA) was performed, followed by a Dunnett’s multiple-comparisons test. Data were considered significant at P < 0.05, with *P < 0.05, **P < 0.01, ***P < 0.001 and ****P < 0.0001.

