## Supplementary material for "tRNA-derived fragments restrict KSHV lytic replication through miRNA-like repression of host pro-viral signalling networks": Supp figs

**Supplementary Figures 1-4 and Table 1.**

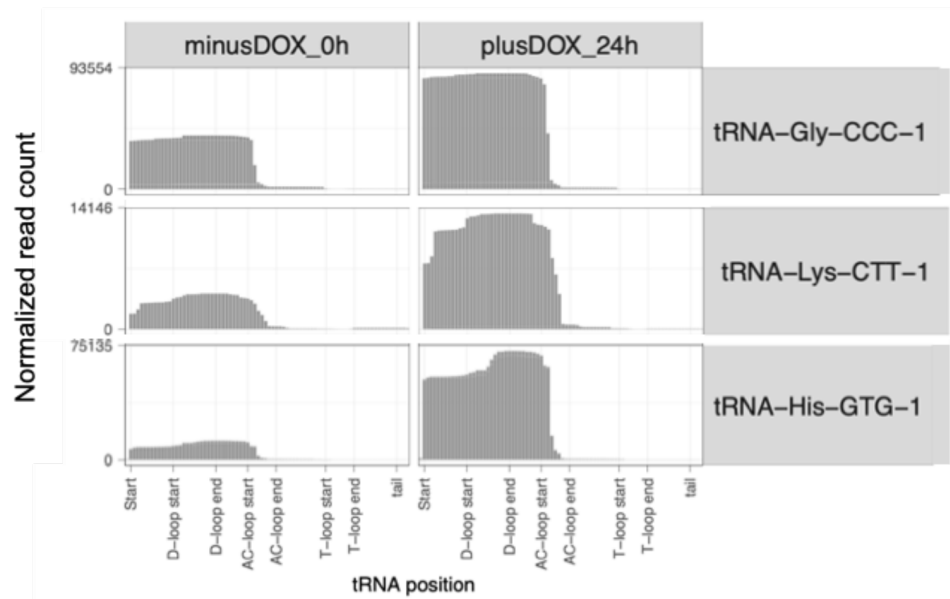

**Fig.S1.** tRF-seq analysis of TREx BCBL1-RTA cells under latent conditions (no DOX) and 24 h after KSHV lytic induction (plus DOX). Data was analysed using tRAX software (https://github.com/UCSC-LoweLab/tRAX) for easy visualisation and characterisation of tDR reads. Reads specifically mapping to the 5' half of the parental tRNAs, tRNA-Gly-CCC-1, tRNA-Lys-CTT-1, and tRNA-His-GTG-1, are more prevalent under lytic conditions <sup>1</sup>.

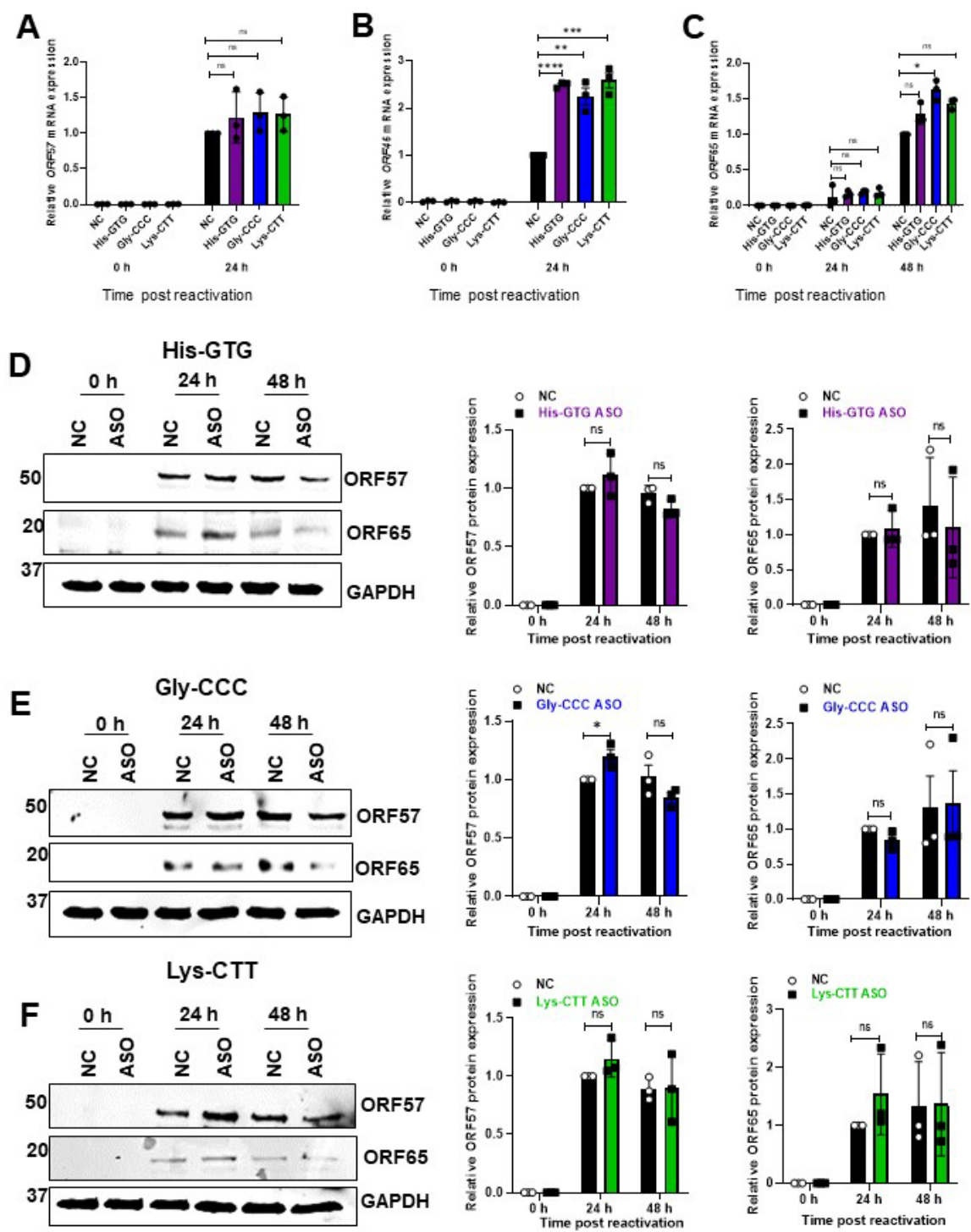

**Fig.S2.** TREx-BCBL1-RTA cells were transfected with a NC inhibitor or the indicated tRF ASOs prior to lytic induction. Samples were harvested at 0 h (latent), 24 h and 48 h post-induction, as specified. RT-qPCR analysis of (A) *ORF57*, (B) *ORF46*, and (C) *ORF65* transcripts normalised to *GAPDH*. Immunoblot analysis of ORF57 and ORF65 protein levels following overexpression of (D) tRF-His-GTG, (E) tRF-Gly-CCC and (F) tRF-Lys-CTT, with densitometric quantification normalised to GAPDH (right). Data are mean ± SD ($n = 3$). Statistical significance was determined by one-way ANOVA (A-C) or unpaired, two-tailed Student's *t* test (D-H), (\*$p < 0.05$, \*\*$p < 0.01$, \*\*\*$p < 0.001$, and \*\*\*\*$p < 0.0001$).

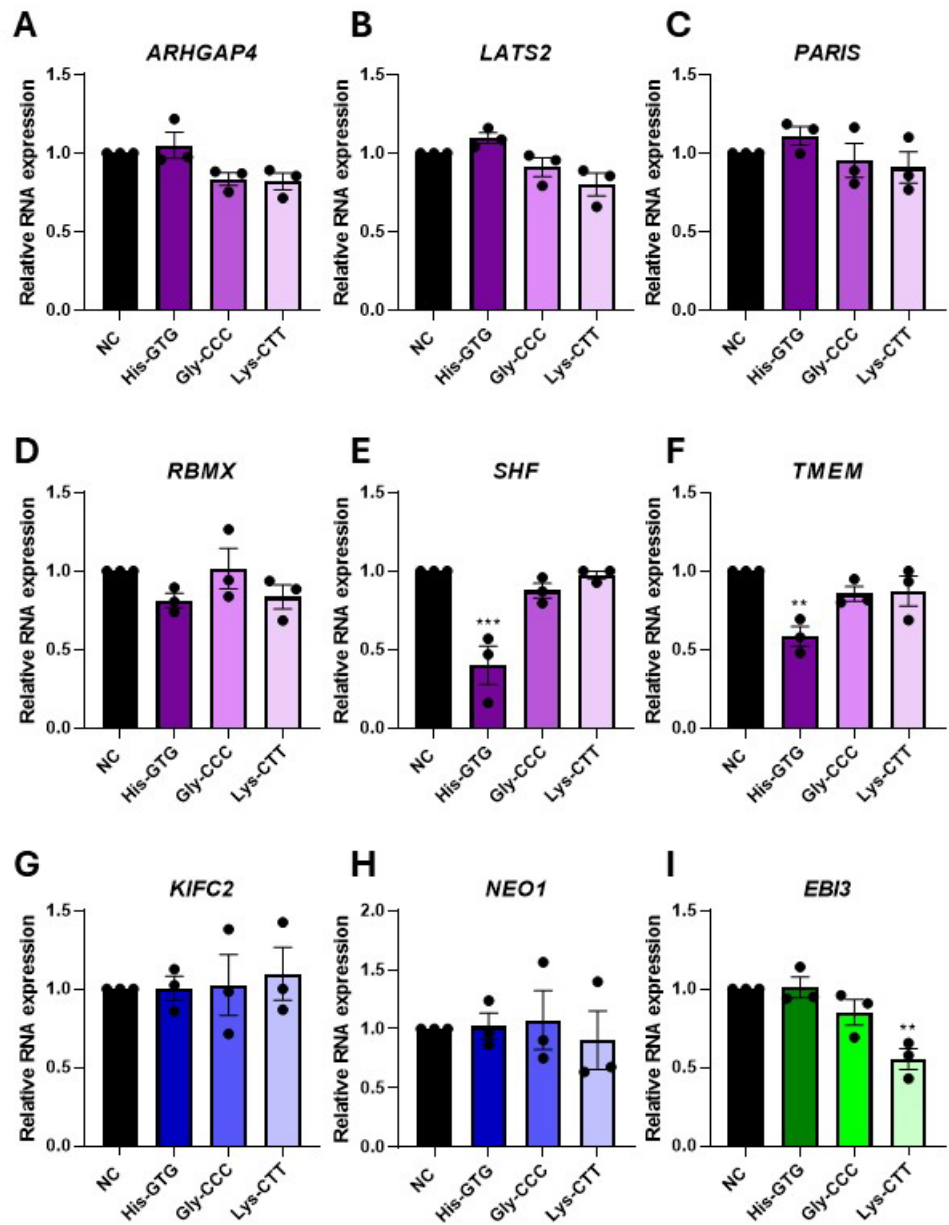

**Fig.S3.** RT-qPCR analysis of predicted targets following tRF overexpression in TREx-BCBL1-RTA cells, normalised to *GAPDH*.

(A-F) Transcript levels of tRF-His-GTG predicted targets.

(G-H) Transcript levels of tRF-Gly-CCC predicted targets *KIFC2* and *NEO1*, respectively.

(I) Transcript levels of the predicted tRF-Lys-CTT target, *EBI3*. Data are mean ± SD (n = 3). Statistical significance was determined by one-way ANOVA, (\*p < 0.05, \*\*p < 0.01, \*\*\*p < 0.001, and \*\*\*\*p < 0.0001).

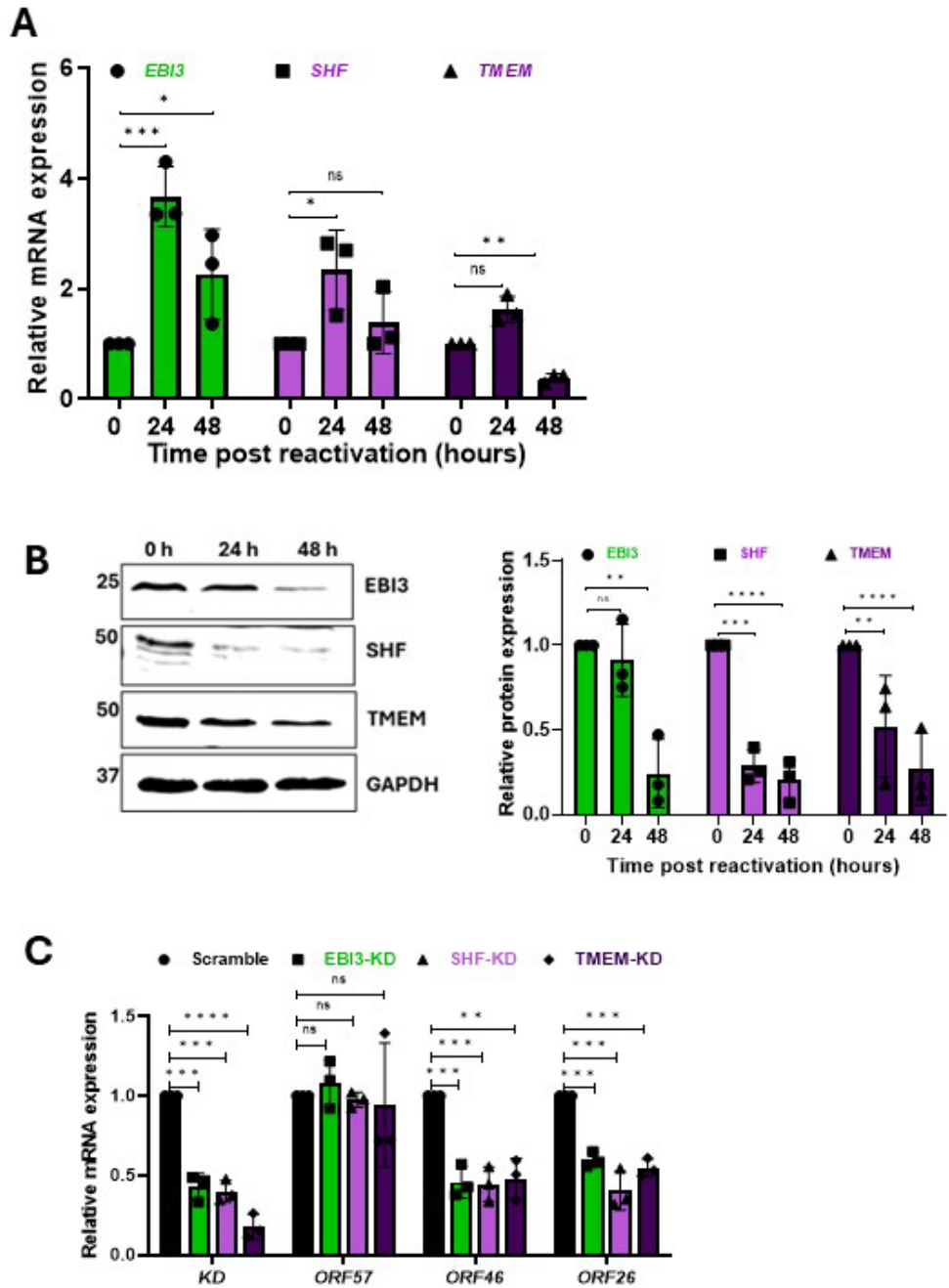

**Fig.S4.** (A) Transcript levels of *EBI3*, *SHF* and *TMEM* in TREx-BCBL1-RTA cells during latency (0 h), or 24 h and 48 h post lytic induction, determined by RT-qPCR and normalised to *GAPDH*.

(B) Corresponding immunoblot analysis of *EBI3*, *SHF* and *TMEM* protein expression, with densitometric quantification normalised to *GAPDH*.

(C) TREx-BCBL1-RTA cells stably expressing shRNAs targeting *EBI3*, *SHF*, *TMEM* (*EBI3*-KD, *SHF*-KD, *TMEM*-KD) or scramble shRNA were analysed 24 h post lytic induction. Knockdown efficiency and expression of *ORF57*, *ORF46* and *ORF26* were assessed by RT-qPCR and normalised to *GAPDH*. Data are mean ± SD ($n = 3$). Statistical significance was determined by one-way ANOVA, (\*$p < 0.05$, \*\*$p < 0.01$, \*\*\*$p < 0.001$, and \*\*\*\*$p < 0.0001$).

**Table S1:** Candidate gene targets of tRF-His-GTG, tRF-Gly-CCC and tRF-Lys-CTT, identified by *tRFtarget*.

| tRF | target gene | Free Energy ($\Delta G$) | complementary length (nt) | Binding region |
| --- | --- | --- | --- | --- |
| His-GTG | ARHGAP4 | -46.9 | 18 | 5'UTR |
|  | FXVD4 | -46.2 | 10 | CDS |
|  | PARIS ZNF746-205 | -46.1 | 11 | 5'UTR |
|  | RBMX | -44.3 | 9 | 5'UTR |
|  | SHF | -52.4 | 25 | 5' UTR |
|  | TMEM237 | -49.2 | 17 | 5'UTR |
| Gly-CCC | AURKAIP1 | -32.1 | 7 | CDS |
|  | KIFC2 | -20.22 | 10 | 3'UTR |
|  | NEO1 | -38.4 | 11 | 3'UTR |
| Lys-CTT | EB13 | -52.5 | 10 | 3'UTR |
|  | GPER108 | -52.9 | 9 | CDS |
